# Capturing and Detecting of Extracellular Vesicles Derived from Single *Escherichia coli* Mother Cells

**DOI:** 10.1101/2023.02.22.529503

**Authors:** Fumiaki Yokoyama, André Kling, Petra S. Dittrich

## Abstract

Cells have a phenotypic heterogeneity even in isogeneic populations. Differences in secretion of substances have been well-investigated with single mammalian cells. However, studies on the heterogeneity of secreted substances at the single-bacterial-cell level are challenging due to the small size, motility, and rapid proliferation of bacterial cells such as *Escherichia coli*. Here, we propose a microfluidic device to achieve an isolated culture of single bacterial cells and capture of extracellular vesicles (EVs) secreted from individuals. The device has winding channels to trap single rod-shaped *E. coli* cells at their entrances. Isolated single mother cells grew constantly up to 24 h, while their daughter cells were removed by flow. The flow carried EVs of the trapped cells along the channel, whose surface was rendered positively charged to electrostatically capture negatively charged EVs, followed by staining with a lipophilic dye to detect EVs by microscopy. Our results underline that the amounts of segregated EVs vary among cells. Moreover, individual responses to perturbation using a membrane-perturbing antibiotic were observed in growth dynamics and EV secretion of living-alone bacteria. The proposed method can be applied to detect other secreted substances of interest, possibly paving the way for elucidating unknown heterogeneities in bacteria.

## 1. Introduction

Cells in all biological kingdom secret extracellular vesicles (EVs),^[1,2]^ which are nano-sized lipid particles containing various biological cargos, such as proteins, nucleic acids, and saccharides.^[3,4]^ EVs have diverse sizes and compositions even when derived from an identical cell population.^[5,6]^ Both eukaryotic and prokaryotic cells possess multiple secretion mechanisms to produce EVs, contributing to the EV heterogeneity.^[4,7]^ It has been revealed that their production and components change in response to environmental fluctuations.^[8,9]^ Moreover, healthy and diseased cells secrete EVs that can differ in size or composition.^[10]^ While secreted from one cell, they can be taken up by other cells in the near or distant environment, thus contributing to cell-to-cell communication.

In recent years, studies on the variation of secreted EVs at the single-cell level, particularly for mammalian cells, have become feasible using different imaging methods^[11]^ as well as microfluidic devices.^[12–16]^ Microcompartments like microchambers and ring-shaped valves were employed to isolate single cells. Subsequent immunostaining allowed the determination of protein cargos such as HSP70 and membrane proteins, such as tetraspanines (CD63, CD81 and others) and receptors such as HER2. These studies have revealed the heterogeneous productivity and cargos of EVs, and enabled the identification of subpopulations of single-cell derived EVs in mammalian cells.

While the EV secretion of mammalian cells has been rigorously studied in bulk and single-cell culture the analysis of EVs secreted from bacterial cells could so far only be shown in bulk culture experiments. The investigation of single bacterial cells such as *Escherichia coli* remained challenging due to their small size, motility, and rapid proliferation. These properties have prevented single-cell compartmentalization as well as the detection of EVs secreted from individual cells. Besides, common molecular markers within bacterial EVs have not yet been identified^[17]^ because bacterial EV cargos are not only heterogeneous with respect to molecular species but also to the secretion levels of the compound present in the EVs.^[3,18]^ In addition, bacterial EVs are also diverse in size and function.^[18,19]^ Despite the difficulty in detecting EVs from bacteria, they have received great attention in recent years because of their role in pathogenesis, antibiotic tolerance, and potential use as vaccine platforms.^[20–22]^ However, our understanding of the heterogeneous EV production in bacteria is still in its infancy due to the technical limitations. Therefore, new approaches are required to keep only single bacterial mother cells confined during culture and detect EVs from the individuals.

In addition to the demands mentioned above, the aspect of single-cell analysis should be highlighted. In microbiology, single-cell analysis by time-lapse microscopy with culture generally means observing and tracking single cells in a growing population or confined in a long channel with their daughters.^[23,24]^ These culture methods of single cells with neighboring daughters cannot entirely rule out cell-to-cell interaction and do not elucidate the lifestyle of living-alone bacteria. Although the most prevalent lifestyle of bacteria in natural environments is thought to be a cellular community known as biofilm,^[25]^ they can also survive in a single-cell state without neighboring cells as unicellular organisms and switch between these unicellular and multicellular states.^[26]^ Therefore, to fully understand bacterial life from unicellular to multicellular aspects, researchers should study not only single bacterial cells in a population, but also isolated single cells, which is also demanding technique for isolated culture of single bacterial cells.

Here, we propose a microfluidic device to achieve an isolated culture of single bacterial mother cells only and detect EVs secreted from the individual cells, inspired by the previous studies using a bacterium-isolating device, the so-called mother machine.^[27,28]^ We detect single cell-derived EVs and linked its production with heterogeneous growth patterns under the perturbation with low concentrations of an antibiotic. This method has potential to open new vistas for elucidating unknown heterogeneities of antibiotic responses and detecting substance secretion from each single bacterial cell.

### 2. Results

### 2.1. Microfluidic Device Enabling Capture of Single *E. coli* Cells

We designed a microfluidic device inspired by a previous study^[27]^ to capture an individual bacterial cell in a narrow channel. In contrast to the former design, our device keeps the initially isolated single-cells only (the “lonely mother cell”), while daughter cells were quickly removed for solitary-cell culture without neighbors (Figure 1A). This modified mother machine device named here isolated mother machine, iMM, enables us to analyze EVs secreted from single bacterial cells over time periods longer than the division time of the cells.

**Figure 1.**
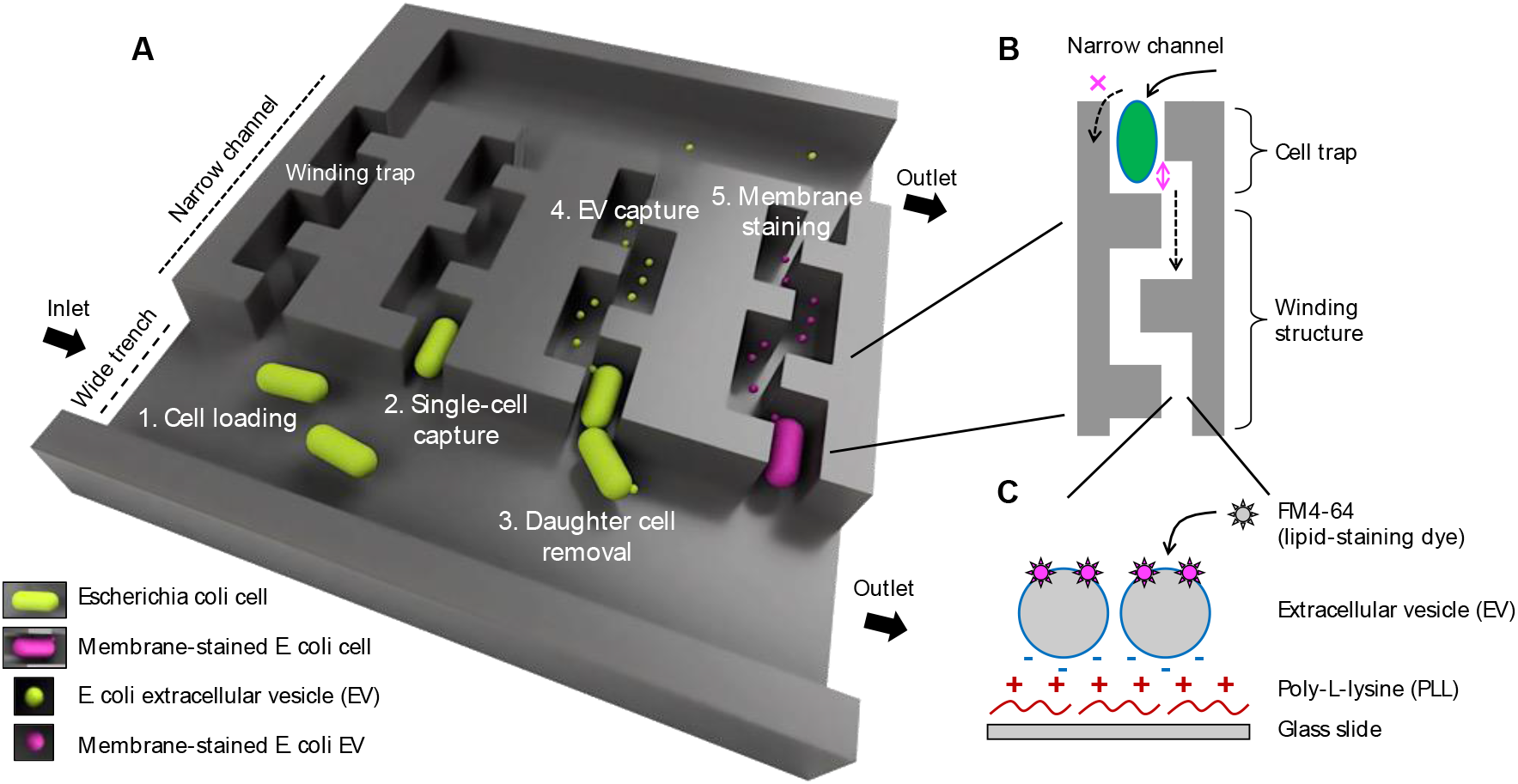
Design of the device and process of EV collection. (A) The microfluidic device enables the isolation of individual bacteria and continuous removal of daughter cells and was therefore named isolated mother machine, iMM, inspired by the original version referred to as “mother machine”. 1. Bacterial cells are loaded into the device through a wide trench by medium flow; 2. A single bacterial cell is captured by a cell trap at the entrance of a narrow channel; 3. During culture, daughter cells from the isolated cell are removed from the cell trap by flow; 4. EVs secreted from the isolated mother cell are carried to the end of a narrow channel by flow and electrostatically captured on the wall; 5. After culture, a lipid-staining dye, FM4-64, is applied to detect labeled EVs by fluorescence microscopy. (B) Top view of a narrow channel. The narrow channel has a 0.8-μm wide corner, and a winding structure restricts the rotation of rod-shaped bacteria like E. coli at the cell trap to prevent its escape, while nano-sized EVs can be carried by flow to the end of the channel. (C) A sketch of electrostatically captured EVs on the device surface from the side view. The glass bottom of the device is rendered positively charged using poly-L-lysine, facilitating negatively charged EVs to attach to the surface by electrostatic forces. A lipid-staining dye, FM4-64, is adsorbed by lipid packing defects of EV membrane and emits fluorescence only at the hydrophobic site like the lipid defects, not or low enough at the aqueous milieu. All the sketches are not to scale.

The iMM device has four inlets/outlets connected to larger channels, referred here to wide trenches (Figure 1 and Figure S1A). The trenches are 100 μm wide and 25 μm high, and are connected to narrow (1.8 µm) and flat (0.85 µm) channels (Figure S1B). The design reported in the literature^[27]^ has a protrusion at the end of narrow channels to prevent isolated cells from escaping but let the flow go through the channel to apply the medium flow efficiently and uniformly, which inspired us to include such protrusions already after 4 μm inside from the entrance of narrow channels to make space for a single *E. coli* cell (Figure S1C). The height of the cell trap was designed to be a bit smaller (0.85 μm) than the *E. coli* cell width reported previously^[29]^ to trap cells robustly at high flow rates that are needed to prevent cell adhesion of the wide trench wall. In addition, the narrow channel following the trapped cell makes a turn (Figure 1B), so that rod-shaped bacteria like *E. coli* cannot tilt or substantially deform to escape the cell trap through the small channel. Furthermore, this cell trap is large enough to capture single cells, but not both mother and daughter cells, and thus the new daughter cells are immediately pushed out of the narrow channel and washed out by flow running in the wide trench.

We made the narrow channel open-end to the wide trench on the other side. This design produces a slow flow through the narrow channels and carry EVs secreted from single mother cells through the winding channel (Figure 1B and S1C). The narrow channels are winding to reduce the flow rate and increase the opportunity of EVs attaching to the wall, whose surface is rendered positively charged with poly-L-lysine to capture negatively charged EVs, followed by lipid staining with a lipophilic dye, *N*-(3-Triethylammoniumpropyl)-4-(6-(4-(Diethylamino) Phenyl) Hexatrienyl) Pyridinium Dibromide (FM4-64) to detect labeled EVs by microscopy (Figure 1C).

Surfaces of the wide trench for medium flow were coated with bovine serum albumin (BSA) to prevent cell adhesion, while those of the narrow channels were coated with poly-L-lysine. This two-pattern modification was achieved by a sequential procedure shown in Figure S2A. First, poly-L-lysine was applied to modify the whole device surface including the narrow channel (Figure S2A Step 1 and 2B). Then, the poly-L-lysine solution was removed from the device by air pressure and incubation at 80 ºC for longer than 16 h (Figure S2A Step 2). Next, BSA solution was applied to modify only the upper-wide trench to mask the positively charged surface by negatively charged BSA (Figure S2A Step 3 and C).^[30]^ In this procedure, the narrow channels are empty because their resistance against aqueous solutions is high enough to prevent them from being filled. As a result, the device surfaces were modified by two patterns: positively and negatively charged surfaces by poly-L-lysine and BSA, respectively, shown using streptavidin-Atto565 and biotin-FITC instead of them (Figure S2D). The poly-L-lysine or BSA coating facilitates or prevents the adhesion of negatively charged substances such as EVs or bacterial cells, respectively.

### 2.2. Growth Dynamics of Isolated Mother Cells of *E. coli*

The microfluidic device was connected with three three-way stop valves via tubing (Figure S3A). The channel design depicted in Figure S3B was important here to apply the cells and medium while preventing contamination in the wide trench and the inlet for medium applying during cell loading. After cell loading, medium was applied to the cells trapped in the narrow channels and collected in the outlets as indicated in Figure S3C. Remaining cells were flushed out by the continuous medium, preventing contamination of the cells growing in the device during culture (Figure S4).

A diluted culture medium with green fluorescence protein (GFP)-expressing *E. coli* at the exponential growth phase was introduced to the coated microfluidic device. Some single cells were isolated by the cell traps (Figure 2A). Under 37 ºC and at 1 mL/h lysogeny broth (LB) flow, the isolated mother cell grew and divided to produce daughter cells with 25 min of doubling time in median (Figure 2B and C). This observation of the single mother cells continued for up to 24 h until we stopped it (Movie S1). We measured the fluorescence intensity of these cells and calculated the cell length and width of each by fitting the fluorescence areas to ellipses, showing their oscillated growth dynamics with mostly 2.5–7 μm in length and almost the same cell width of about 1.2 μm during culture (Figure 2D and Figure S5A and Figure 2E and Figure S5B, respectively). Occasionally, a cell in this measurement stopped growing and expanded its cell width around the end of the measurement (Figure S5 and Movie S2). The measurement for cell length means that 33% of cell length of 6-μm elongated mother cells was at the outside of the 4-μm cell trap, and EVs secreted from the outer body part were excluded from the analysis in this method. Some cells, in particular abnormally elongated cells, were pushed out by the flow outside of the cell trap and detached at some points.

**Figure 2.**
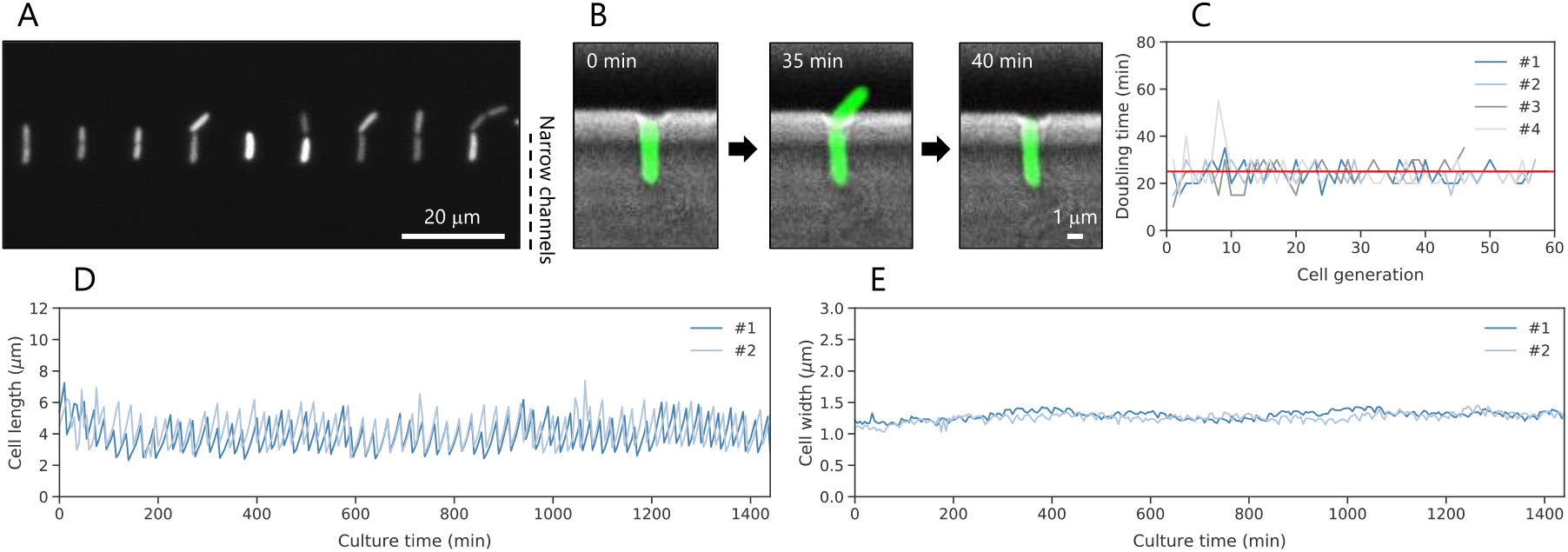
Growth dynamics of isolated mother cells of *E. coli.* (A) A representative image of GFP-expressing E. coli mother cells isolated in cell traps. (B) Growth and daughter cell removal of an isolated mother E. coli cell. A fluorescence image of a GFP-expressing cell and a brightfield image of a narrow channel are overlayed. The daughter cell is clearly visible 35 min after cell isolation, and was removed by medium at t = 40 min. (C) Doubling time of isolated single cells. The doubling time was calculated from the fluorescence images of the cells. The red line represents the median doubling time of the data (25 min). (D and E) Physical parameters obtained from the fluorescence images of single-cell growth. The growth of isolated single cells was monitored for up to 24 h, and their cell length (D) and width (E) were measured from the fluorescence images. Each data line indicated single bacterial cells. Two represented data are shown here, and the two others are in Figure S5. All images were taken by multiple independent experiments.

In addition, daughter cells sometimes attached to the wall of narrow channels after cell division and prevented the single-cell analysis. These data were removed from the analysis. The obtained growth patterns were similar to previous studies using the original *mother machine*,^[28,31]^ indicating the trapped cells in our modified design grew well.

### 2.3. Heterogeneous Growth Dynamics of Single Isolated Mother Cells of *E. coli* under Antibiotic Exposure

To investigate the antibiotic effects on EV production at the single-bacterial-cell level, we used a membrane-perturbing antibiotic for Gram-negative bacteria, polymyxin B. We selected this compound because it has been reported that 750 ng/mL of polymyxin B induces EV production in the bulk culture of *E. coli* as part of an immediate defense strategy.^[32]^ Growth curves of *E. coli* confirm that cells grew well in static culture with a multi-well plate when the polymyxin B concentration was less than 500 ng/mL (Figure S6A). Images taken with an agarose pad showed that cells at the exponential growth phase had similar morphology under 0 and 250 ng/mL of polymyxin B (Figure S6B). The cell size measured from these fluorescence images showed that the median cell length at 250 ng/mL of polymyxin B (4.0 μm) was slightly larger than that without polymyxin B (3.6 μm) (Figure S6C). The median cell width at 250 ng/mL of polymyxin B (0.94 μm) did not show significant differences from that without polymyxin B (0.95 μm) (Figure S6D), meaning that captured *E. coli* cells were expected not to escape from the cell trap because they do not shrink or elongate too much even during culture in the presence of polymyxin B at 250 ng/mL. The population smaller than 0.85 μm in cell width at the exponential growth phase without polymyxin B, which was expected to be captured by the 0.85-μm high cell trap, was approximately 15% (Figure S6D).

We measured the growth of single mother *E. coli* cells on chip in the presence of polymyxin B (Figure 3A). Although most cells in the absence of polymyxin B grew well for 24 h until we stopped the observation, they elongated at some points due to the division deficiency as mentioned above (Figure 3A top, Figure S7 top, and Movie S3). On the other hand, in the presence of polymyxin B at 50 and 250 ng/mL, we could not observe the cells grew over the full 24h (Figure 3A middle and bottom, Figure S7 middle and bottom, and Movie S4 and S5, respectively). Under these conditions, we observe the gradual loss of GFP fluorescence (Movie S5). The dead cell, however, was still captured, as we could stain the membrane with FM4-64 after 24-h culture (Figure S8), and therefore the killing mechanism by polymyxin B did not cause the complete membrane fragmentation of lysed cells, and most of their membranes were not included in the following EV analysis by lipid staining. The cell length at each time point was plotted and showed the median cell length was reduced for higher polymyxin B concentration (4.0, 3.9, and 3.6 μm at 0, 50, and 250 ng/mL of polymyxin B, respectively, Figure 3B). The median doubling time at all concentrations of polymyxin B was 25 min (Figure 3C).

**Figure 3.**
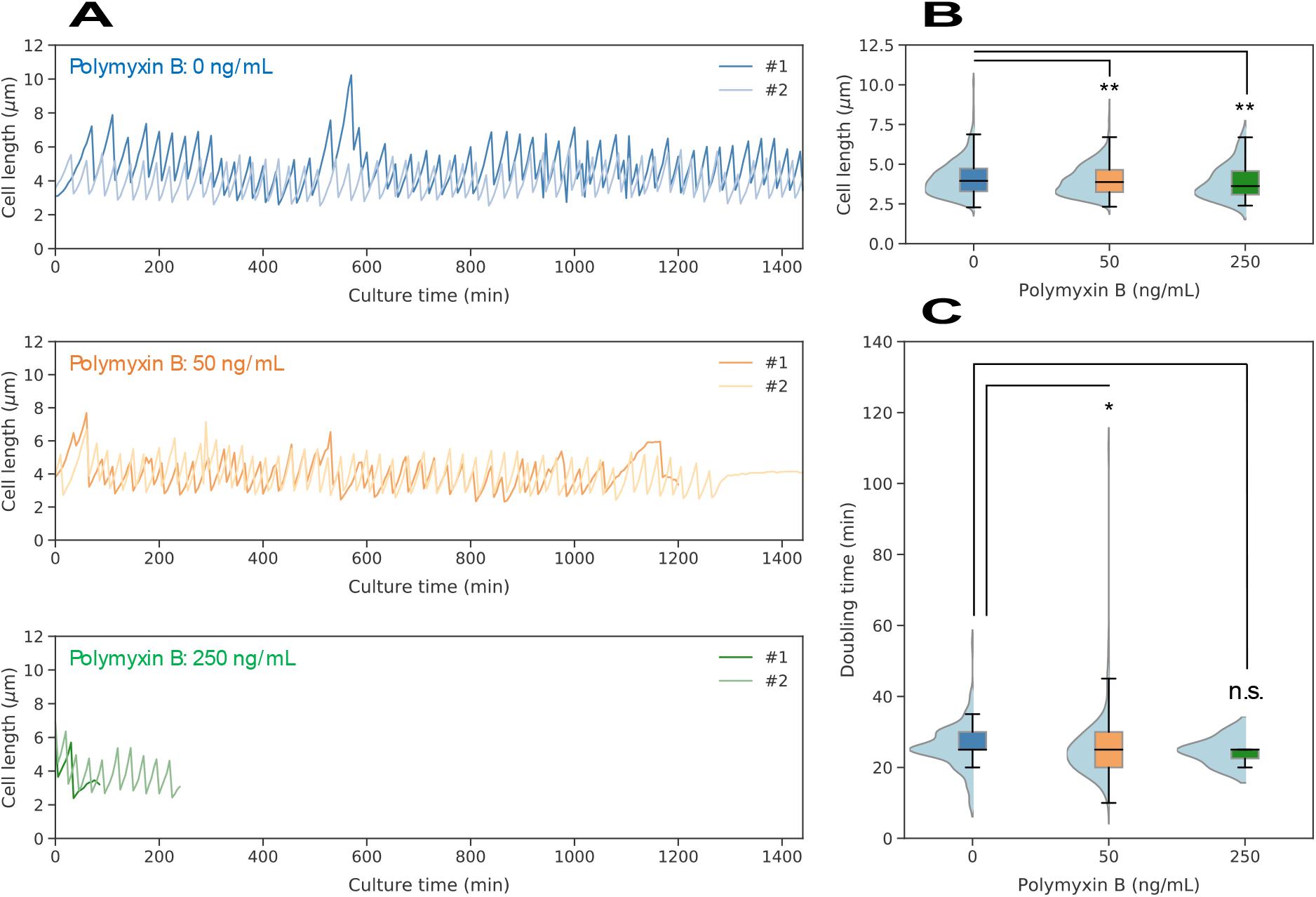
Heterogeneous growth of single *E. coli* cells under antibiotic exposure. (A) Time course of cell length dynamics at different concentrations of polymyxin B. The growth of captured single cells was monitored for up to 24 h. Each data line represents a single bacterial cell (additional examples in Figure S7). (B) Cell length of each mother cell obtained from cell length dynamics at different concentrations of polymyxin B. (C) Doubling time of each mother cell obtained from the time course of growth dynamics. Each black line within the boxes represents the median, each lower and upper edges of the boxes represent the 25 and 75 percentiles of the data, respectively, and each error bar represents the maximum and minimum values of the box plots (n > 84 in B and 11 in C). The gray dots on the right side of the boxes represent each data point. The light blue graphs on the left side of the boxes represent each data distribution. *p < 0.01, **p < 0.05, and n.s.: no significant difference (two-sided Brunner-Munzel test).

### 2.4. Detection of *E. coli* Extracellular Vesicles on Microfluidic Device

To capture EVs on the microfluidic device, we used electrostatic force between EVs and the coated glass surface. The zeta potential measurement showed that EVs and poly-L-lyisne had -12.7 and 35.8 mV in the median, respectively (Figure 4A), indicating that poly-L-lysine-coated surface can electrostatically capture EVs. To test this, we modified the polystyrene surface of a multi-well plate using poly-L-lysine, then applied EVs, followed by lipid staining with a lipophilic dye, FM4-64. The median value of FM4-64 fluorescence intensity for EV lipid membrane was significantly higher on the coated surface (87.0 a.u.) than those on the non-coated surface (43.0 a.u.) and using EV-free supernatant instead of EV suspension on the coated surface (17.0 a.u.) (Figure 4B). This result indicated that the combination of electrostatic force and lipid staining was applicable for EV detection in the narrow channels of the proposed microfluidic device.

**Figure 4.**
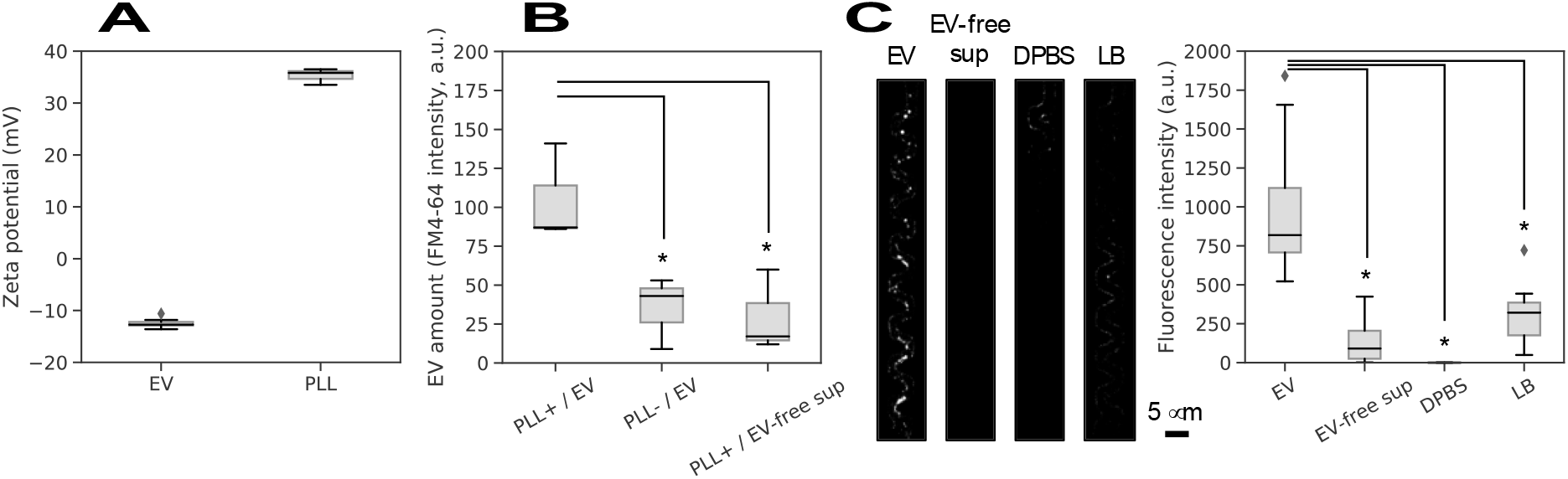
Detection of extracellular vesicles in narrow channels of microfluidic device. EV capturing and detection by electrostatic interaction and lipid staining. (A) Zeta potential of EVs and poly-L-lysine (PLL) in DPBS. (B) EV detection on a 386-well plate with polystyrene surface modified with and without poly-L-lysine. As a control, EV-free supernatant (sup) or a non-coated plate were used. (C) Representative images of detected EVs on the microfluidic device by lipid-staining with FM4-64 (left) and the fluorescence intensity of EVs, EV-free sup, DPBS, LB quantified with the images (right). Each black line within the boxes represents the median, each lower and upper edges of the boxes represent the 25 and 75 percentiles of the data, respectively, and each error bar represents the maximum and minimum values of the box plots (n = 3 or 9 in Figure 4A and B, n = 18 in Figure C). The gray dots on the right side of the boxes represent each data point. *p < 0.01 (two-sided Brunner-Munzel test). All images were taken by multiple independent experiments.

Next, purified EVs were introduced into the coated device. After incubation to allow EVs being captured by the poly-L-lysine-coated surface, we applied FM4-64 to stain the membrane of the captured EVs. We observed fluorescent spots with diameters < 1 µm on devices incubated with EVs, which we did not observe when the device was incubated with EV-free supernatant, Dulbecco’s phosphate-buffered saline (DPBS), or LB (Figure 4C left). Comparison of the fluorescence intensity in a region of interest along the narrow channels confirmed the difference for the channel exposed to EV suspension from that to the controls (Figure 4C right).

### 2.5. Quantification of EVs Derived from Single Mother Cells of *E. coli* under Antibiotic Exposure

Finally, we collected the EVs from individual cells and cells under antibiotic treatment. At the cell trap, isolated mother *E. coli* cells divided and produced EVs by budding during culture (Figure 5A). After 24-h culture for all the conditions, FM4-64 was applied to the microfluidic device to stain lipid membranes of the mother cells and the captured EVs in the narrow channels. The winding structures, excluding cell traps of the narrow channels, were set as a region of interest (ROI, Figure 5B). The microscope images (Figure 5C) indicated that we could clearly harvest and detect EVs in the winding channels, where the cell trap was occupied by a bacterial cell. Untreated cells segregated visible more EVs after 24 hours, while we observe less EV signals for treated cells. However, for treatment with the larger concentration of polymyxin B, the cells grew only a short time. The dead cells remained in the trap what we confirmed after 24 hours, however, we assume that EV production discontinued when the cell growth and division stopped. Therefore, we plotted the EV production from individual *E. coli* cells (i.e. the integrated fluorescence signal) as a function of total growing time (Figure 5D). The values of the EV intensity varied from cell to cell, even within the same culture condition and we observed low EV numbers in several cases. We often found aggregated and chain-like structures, confirmed in bulk measurements (Figure S9 and Movie S6), which hindered enumeration of EVs and were taken out from analysis. In future, reliable staining procedures may support a more robust quantification of the EV production.

**Figure 5.**
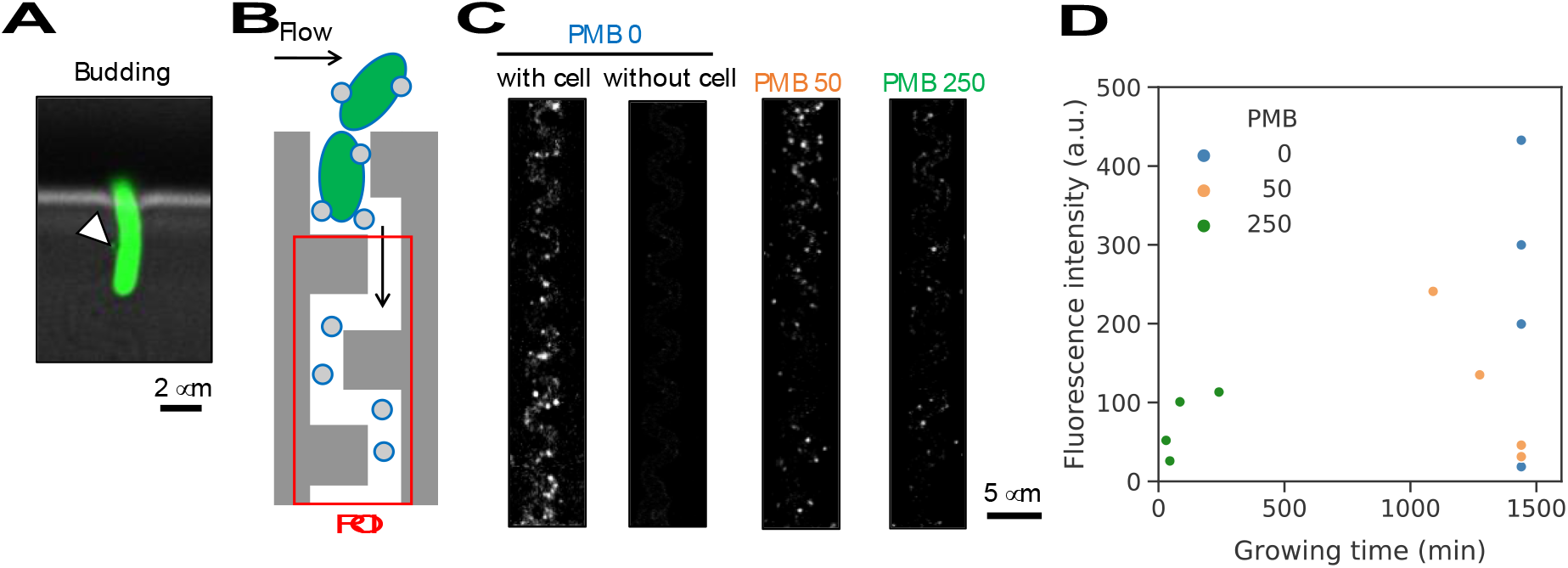
Extracellular vesicle secretion of single *E. coli* cells. (A) A representative image of an E. coli cell with membrane budding. A fluorescence image of a GFP-expressing cell and a brightfield image of a narrow channel are overlayed. Green: GFP-expressing E. coli cell, and white arrowhead: budding membrane. (B) A sketch of single mother cell-capturing narrow channel and region of interest (ROI), shown in red. Green rod shape: E. coli cell, and gray sphere: EV. The sketch is not to scale. (C) Representative images of captured EVs after 24-h culture at different concentrations of a membrane-perturbing antibiotic to Gram-negative bacteria, polymyxin B. PMB 0, 50, and 250 represents polymyxin B at 0, 50, and 250 ng/mL, respectively. White dot: EVs stained with FM4-64. (D) Individual fluorescence intensity of lipid-stained EVs as a function of growing time from single mother cells at different concentrations of polymyxin B.

We compared these observations with results from bulk culture, where EVs were collected after 16 h, when the cells were already in the stationary growth phase. Here, we found that cells under 250 ng/mL of polymyxin B secreted bigger EVs (192 nm in median diameter) compared with those at 0 and 50 ng/mL of the antibiotic (104 and 107 nm, respectively) (Figure S10A and B). Polymyxin B at 250 ng/mL also induced a larger EV production (Figure S10C and D). The induced EV production by polymyxin B at 250 ng/mL was thought not to attribute to the larger number of cells in the cultured medium at the stationary phase because these low concentrations of the antibiotic did not affect the growth curves in the static culture, and the OD600 values between different polymyxin B concentrations did not show significant differences in the bulk culture (Figure S6A and S10E, respectively). Our single-cell data correlate with the bulk measurements, if we consider the short growth time for cells treated with polymyxing at 250 ng/mL. By dividing the fluorescence signal by the growing time (Figure S11), a high vesiculation under the stronger antibiotic exposure is determined, whose tendency is also in accordance with our data in the bulk culture (Figure S10C and D) and a previous study by Manning and Kuehn (26).

## 3. Conclusion

In conclusion, the proposed iMM device has enabled solitary-cell culture by tracking single bacterial cells without surrounding cells by removing their daughter cells by flow for up to 24 h, to assess growth rates, morphological parameters, and substance secretion of the individuals. In addition, this device easily enables medium exchange, enabling us to analyze the responses of living-alone cells to environmental fluctuations. This unique feature of the device potentially reveals bacterial responses and heterogeneities in given environments as a unicellular organism, extending our knowledge of bacterial life in unicellular states.

To investigate the effects of environmental perturbation, we added low concentrations of a membrane-perturbing antibiotic to Gram-negative bacteria, polymyxin B, to medium and measured the growth dynamics of each cell. Although cells in 200-μL bulk culture of a multi-well plate statically grew well under less than 500 ng/mL of polymyxin B (Figure S6A), living-alone mother cells without neighbors died in the presence of polymyxin B at 50 and 250 ng/mL after around 1200 and 200 min of culture time, respectively (Figure 3A and Figure S7). During this time, we could collect EVs from individual cells and found a high diversity. A particular challenge was the clustering of EVs, confirmed by bulk measurements (Figure S9), which prevented reliable quantification with the number of EVs secreted from single cells. As a preliminary result, we find a relatively large secretion of EVs in the isolated culture of single bacterial cells under 250 ng/mL of polymyxin B treatment, before they stop growing and dividing (Figure S11). This tendency was seconded by measurements in bulk culture (Figure S10C and D), raising the question how the cells, stressed by antibiotic compounds, increase the vesiculogenesis without growth defect in a population and whether this is involved in the emergence of resistance as a collective behavior.^[33]^ Future work will include improving throughput by optimizing the conditions for bacterial cell capture and preventing cell escape to implement accurate statistics and reveal more details of EV heterogeneity at the single-bacterial-cell level.

The unique advantage of the iMM device can be extended in the detection of other secreted substances derived from single bacterial cells. In principle, the glass surface of the narrow channel on the proposed device can be modified using several different types of antibodies against molecules of interest with established coating methods for microfluidic devices, allowing multiplexing.^[12–16]^ This flexibility of the device extends its applicability to other secreted molecules, like secreted proteins, from single bacterial cells. Bacteria have several types of secretion mechanisms,^[34,35]^ and some secreted proteins possess toxicity and are related to pathogenesis.^[36]^ In general, such secreted proteins have been analyzed using culture supernatants of bulk culture, showing the results from the averaged analysis. Therefore, the proposed iMM device has the potential to elucidate deeper characteristics in protein secretion at the single-bacterial-cell level in an isogenic cellular population. Overall, the iMM device and the proposed solitary-cell culture method potentially extend our knowledge of unknown bacterial heterogeneities and living-alone unicellular states.

## 4. Experimental Section

### Bacterial Strains and Culture Conditions

For monitoring bacterial growth, *E. coli* K-12 MG1655 (pSEVA271-*sfgfp*) was used.^[37]^ A glycerol stock of this *E. coli* was streaked onto an LB plate containing 1.5% agar and incubated at 37 ºC for 24 h and stored at 4 ºC until used. A single colony on the plate was inoculated in LB with kanamycin (50 μg/mL) and cultured at 200 rpm and 37 ºC for 16 h using an Ecotron incubator (Infors, Bottmingen, Switzerland). Ten microliters of the pre-cultured medium were subcultured in 1 mL of LB with Kanamycin (50 μg/mL) and cultured at 200 rpm and 37 ºC approximately for 2 h until cells entered the exponential growth phase (the optical density at 600 nm, OD600, reached around 0.3). This cultured medium was used for the single-cell culture in the microfluidic device. For the bulk culture, 40 μL of the pre-cultured medium was subcultured in 4 mL of LB with kanamycin (50 μg/mL) and cultured at 200 rpm and 37 ºC for 16 h until cells entered the stationary growth phase (OD600 values reached around 7.0). The values of OD600 were measured using a NanoPhotometer spectrophotometer (Implen, Munich, Germany). A membrane-perturbing antibiotic for Gram-negative bacteria, polymyxin B (Sigma-Aldrich, St. Louis, MO, United States), was used to investigate the stress responses of each cell in terms of EV secretion when needed.

### Microfluidic Device Fabrication

The dimethylpolysiloxane (PDMS) microfluidic devices were fabricated using a 10:1 mass ratio of Sylgard 184 Silicon Elastomer Base and Curing Agent (Dowsil, Midland, MI, United States). These reagents were mixed well using a plastic spatula and degassed using a vacuum pump until visually bubble-free. Approximate 5 g of the mixture was poured onto the master mold and placed at 80 ºC for 3 h. After curing, the fixed PDMS was peeled off from the master mold and unnecessary parts of the PDMS block were cut off. Four inlets/outlets were punched using a 1.5-mm biopsy puncher (Integra LifeSciences, Princeton, NJ, United States). The surface of the device was cleaned using adhesive tape. A No. 1.5 microscopy glass slide (Biosystems, Muttenz, Switzerland) was cleaned by washing with acetone, isopropanol, and water, then dried using nitrogen gas and a heater at 150 ºC. The device and the glass slide were plasma-activated using a PDC-32G plasma cleaner (Harrick Plasma, Ithaca, NY, United States) at less than 0.9 mbar for approximately 1 min (18 W) and bound together. The glass-bounded device was put on a heater at 150 ºC for 5 min and stored at room temperature until used.

### Two-Patterned Surface Functionalization of Microfluidic Device

0.01% (0.1 mg/mL) poly-L-lysine (70,000–150,000 molecular weight, Sigma-Aldrich) was applied to a microfluidic device using a polytetrafluoroethylene (PTFE) tube (inner diameter 0.25 mm, outer diameter 1.59 mm, BGB, Orsa, Sweden, or PKM Konrad, Rotkreuz ZG, Switzerland) and 1 mL syringe (Becton, Dickinson and Company, Franklin Lakes, NJ, United States) from the lower-left inlet (Figure S1A and S2A Step 1), and incubated at 20 ºC for 1 h. Then, the air pressure was applied from the same inlet using a PTFE tube and a syringe to remove the solution from the device (Figure S2A Step 2). The device was heated at 80 ºC 19 h to evaporate the remaining poly-L-lysine solution. A top wide trench of the device was refilled with 10 μL of BSA by pipetting to the upper-left inlet, followed by flushing the solution with 20 μL of DPBS (KCl 0.2 g/L, KH_2_PO_4_ 0.2 g/L, NaCl 8 g, Na_2_HPO_4_ 1.15 g/L, pH = 7.0–7.3, Thermo Fisher Scientific, Waltham, MA, United States) (Figure S2A Step 3). The narrow channels of the device were filled with LB using a PTFE tube and a syringe at the lower-left inlet and stored at 20 ºC until use (less than 2 h) (Figure S2A Step 4). The poly-L-lysine coating lenders the surface of the device positively charged to electrostatically capture negatively charged EVs, and the BSA-coated surface prevents cell adhesion to the wall of a wide trench. For the visualization of the narrow channels and the wide trenched filled with the solution, 5 μg/mL streptavidin-Atto565 and 5 μg/mL biotin-fluorescein isothiocyanate (FITC) were used instead of poly-L-lysine and BSA at Step 1 and 3, respectively (Figure S2B and C). Brightfield and confocal fluorescence images were taken with ×20 objective (0.75 numerical aperture), as described below, for FITC (emitter: ET525/50m at 525/50 nm, Chroma Technology, Olching, Germany) using 5% laser power of 488 nm and 50 ms exposure time and for Atto565 (emitter: FF01-609/54 at 609/54 nm, Semrock, DEX Health & Science, LLC Rochester, NY, USA) using 50% laser power of 561 nm and 1 s exposure time. For the visualization of narrow channels in Figure S2D, the background of the overlaid image between Figure S2B and C was subtracted with a 50-pixel rolling ball radius in ImageJ/Fiji version 2.3.0/1.53f, then adjusted in its brightness.

### Time-Lapse Microscopy for Growth Measurement

Cells in the exponential growth phase were pelleted by centrifugation at 6800 × g and 20 ºC for 5 min and washed in DPBS twice. One-thousand-time diluted cell suspension in a fresh medium was applied to the upper-left inlet with 100 μL/min flow rate, while fresh medium with kanamycin (50 μg/mL) and polymyxin B, when needed, was applied from the lower-right inlet with 120 μL/min flow rate to keep the inlet non-contaminated (Figure S3B). Once some cell traps were occupied by single cells for approximately 5 min, the flow containing cells was stopped, the three-way stop valve at the upper-left inlet for cell suspension was open to the outlet for waste, and the upper-right outlet used for waste in cell loading was closed with a three-way stop valve. This process allowed the fresh medium flow to direct to the narrow channels and then to the lower-left outlet (Figure S3C). The flow rate of the medium was set at 1 mL/h for the growth measurement. Fluorescence images before and after valve opening and closing were taken by epi-fluorescence microscopy for excitation and optical filters and dichroic mirrors for GFP, as described right below, using 1 s exposure time to show the flow direction in Figure S3.

The trapped cells were imaged in brightfield mode of a Nikon ECLIPSE Ti 2 inverted microscope (Nikon, Tokyo, Japan) with a light-emitting diode (LED) illumination system (CoolLED, Andover, UK), a motorized stage, ×100 objective (1.49 numerical aperture, with immersion oil), a Hamamatsu Orca Flash 4.0 V2 complementary metal-oxide-semiconductor (CMOS) camera with a sensor size of 2044×2048 pixels (Hamamatsu Photonics, Shizuoka, Japan). Fluorescence images of the cell traps were taken by epi-fluorescence microscopy mode using a Spectra X LED unit (Lumencor, Beaverton OR, United States) for excitation and optical filters and dichroic mirrors for GFP (exciter: ET470/40x at 470/40 nm, emitter: ET525/50 at 525/50 nm, dichroic: T496lpxr, Chroma Technology) using 50 ms exposure time every 5 min for 24 h. Imaging was controlled by Visitron VisiView.

For the acquisition of cell morphological parameters, the images were processed using ImageJ/Fiji version 2.3.0/1.53f. First, the scale was set using the known scale provided by Visitron VisiView. The background was subtracted with a 50-pixel rolling ball radius. Then, the threshold was automatically adjusted for the dark background with the default setting. By particle analysis, the fluorescence cells were fit to ellipses, and their major and minor length was measured as cell length and width, respectively. The doubling time was calculated from the cell length data.

### Purification of Extracellular Vesicles by Ultracentrifugation

EVs secreted from *E. coli* cells were collected from cultures at the exponential and the stationary growth phase. The cells were pelleted by centrifugation at 6,800 × g and 20 ºC for 10 min. The supernatant was centrifuged at 13,000 × g and 20 ºC for 15 min to remove the remaining bacterial cells. The supernatant was filtered through a 0.2-µm pore polyethersulfone (PES) filter to remove the remaining debris. EVs were obtained by ultracentrifugation of the filtrate at 100,000 × g (average centrifugal force) and 4 ºC for 2 h with a centrifuge (SORVALL WX Ultra Series, Thermo Electron Corporation, Waltham, MA, United States). The pellets were ten times concentrated by resuspending in DPBS and used as EVs. The supernatants without EVs after ultracentrifugation were used as EV-free supernatants. These EVs and EV-free supernatants were kept at 4 ºC for less than two days.

### Zeta Potential of Extracellular Vesicles and Poly-L-Lysine

The zeta potential of EVs in DPBS and 0.01% poly-L-lysine (Sigma-Aldrich) was measured at 20 °C using a Zetasizer Nano dynamic light scattering analyzer and Disposable Folded Capillary Cell DTS1070 (Malvern Panalytical, Worcestershire, UK). The distribution data were obtained by repeating up to 100 cycles.

### On-Plate Assay for Extracellular Vesicle Detection

The surface of a 384-well black plate (Greiner, Kremsmünster, Austria) was incubated with 20 μL of 0.01% poly-L-lysine (Sigma-Aldrich) at 20 °C for 1 h, followed by washing with 50 μL of DPBS (Thermo Fisher Scientific) twice. Twenty microliters of purified EVs or EV-free supernatant were applied to the wells and incubated at 37 °C for 24 h, followed by lipid-staining with 20 μL of 5 μg/mL FM4-64 (Thermo Fisher Scientific) at 20 °C for 30 min in the dark. The fluorescence intensity of FM4-64 was measured at 515/20 nm excitation and 635/20 nm emission wavelength using Cytation 5 Cell Imaging Multi-Mode Reader (BioTek, Winooski, VT, United States). A non-coated plate was used with EVs as a control.

### Extracellular Vesicle Detection by Microscopy with Lipid Staining

After culture of cells in the microfluidic device for 24 h or applying purified EVs, 1 μg/mL FM4-64 (Thermo Fisher Scientific) in DPBS were applied to the device by flow at 1 mL/h for 30 min in the dark. Images of the narrow channels were taken using a Nikon ECLIPSE Ti 2 inverted microscope (Nikon) with a motorized stage, ×100 objective (1.49 numerical aperture, with immersion oil), a Hamamatsu Orca Flash 4.0 V2 CMOS camera with a sensor size of 2044×2048 pixels (Hamamatsu Photonics). Fluorescence images were taken at the bottom of the device by confocal microscopy mode for FM4-64 (emitter: FF01-609/54 at 609/54 nm, Semrock) using 50% laser power of 561 nm and 1 s exposure time. VS Laser Modul 1865 with Laser-Merge and VS Phase switcher for excitation at 561 nm was used for the laser illumination. Imaging was controlled by Visitron VisiView.

For the detection of EVs on an image, the images were processed using ImageJ/Fiji version 2.3.0/1.53f. The brightness and contrast were adjusted, and each narrow channel excluding a cell trap region was selected by a rectangle as a region of interest (ROI), shown in Figure 5B. We integrated the fluorescence signals of each ROI and used the value as a means of EV number.

## Supporting Information

Supporting Information is available from the Wiley Online Library or from the author.

## Acknowledgements

We thank Dr. Voichita Mihali and Prof. Cornelia G. Palivan (University of Basel) for nanoparticle tracking analysis, Dr. Jonas M. Nikoloff (ETH Zurich) for help with microscopy, the team of Single-Cell Facility and of Clean Room Facility of the Department of Biosystems Science and Engineering (ETH Zurich) for their valuable help and advice, and Prof. David Juncker (McGill University) for the advice on using lipid-staining for EV detection. We gratefully acknowledge the funding from the European Research Council (ERC Consolidator Grant No. 681587 to P.S.D.), and the National Centre of Competence in Research (NCCR) AntiResist funded by the Swiss National Science Foundation (grant number 51NF40_180541).

## Author Contributions

F.Y. designed the research, performed all experiments, and analyzed the data. A.K. fabricated master molds. P.S.D. supervised the work. F.Y and P.S.D. discussed the results and wrote the paper. All authors reviewed, commented the manuscript, and approved the final version of the manuscript.

## Data Availability Statement

The data that support the findings of this study are available from the corresponding author upon request.

