## Supplemental information for "Capturing and Detecting of Extracellular Vesicles Derived from Single *Escherichia coli* Mother Cells"

ETH Zurich

Basel 4058, Switzerland

F. Yokoyama

Current address: Department of Physics

The University of Tokyo

Tokyo 113-0033, Japan

**Supporting Experimental Section**

*Master Mold Fabrication.* The master molds for the PDMS chip replication were fabricated in a cleanroom environment. The mold designs were enlarged to 101.16% of their designated size to account for the shrinkage of the cured PDMS. First, a 4-inch silicon wafer was plasma treated and subsequently spin coated with a layer of GM1030 (Gersteltec, Switzerland) photoresist at 1250 rpm to achieve a height of 0.85 µm. The wafers were softbaked at 95°C for 2 min and exposed with UV light (160 mJ/cm$^2$, i-line) through a chrome mask (Selba, Versoix, Switzerland) incorporating the narrow channel structures using a MA-8-mask aligner (Süss MicroTec, Munich, Germany). After a post-exposure bake for 2 min at 95°C another SU-8 3025 photoresist (Kayaku Advanced Materials, Westborough, MA, United States) layer was spin-coated at 4000 rpm to result in a final thickness of 25 µm. The resin was again soft-baked for 15 min at 95°C and UV-exposed (160 mJ/cm$^2$, i-line) through a chrome mask to define the wide channel structures of the molds. After a post-exposure bake for 3.5 min at 95°C the master molds were developed for 6 min in a mr-Dev 600 (micro resist technoglogy, Germany) developer bath. Finally, the structures were hard-baked at 120 °C for 45 min before being treated with trichloro(1H,1H,2H,2H-perfluorooctyl)silane (Sigma Aldrich, United States).

*Laminar Flow to Prevent Contamination of Narrow Channels.* Cells were applied from the upper-right inlet with 1 µL/min, while DPBS was applied to the lower-right inlet with 50 µL/min. After 1 min, the upper-right inlet was closed, and the flow rate of the medium was set 1 mL/h, the same as the growth measurement. Fluorescence images were taken by epi-fluorescence microscopy mode for GFP, as described in Experimental Section, using ×20 objective (0.75 numerical aperture) and 500 ms exposure time.

*Growth Measurement Using Plate Reader.* Two microliters of the precultured medium were subcultured into 200 μL of LB with kanamycin (50 μg/mL) in a well of a 96-well plate (Greiner). This plate was incubated at 37 °C, and $OD_{600}$ values were measured using Cytation 5 Cell Imaging Multi-Mode Reader (BioTek) every 30 min for 24 h.

*Morphological Analysis and Cell Size Measurement by Fluorescence Microscopy.* Cells in the cultured medium were pelleted by centrifugation at 6,800 × g and 20 ºC for 5 min and then resuspended in DPBSS. This procedure was repeated twice. Two microliters of the suspension were loaded on No. 1.5 microscopy glass slide (Biosystems) and covered with 1% agarose pad. Fluorescence images of the cells were taken by confocal microscopy mode for GFP, as described in Experimental Section.

For the acquisition of physical cell parameters, the images were processed using ImageJ/Fiji version 2.3.0/1.53f. First, the scale was set using the known scale provided by Visitron VisiView. The background was subtracted with a 50-pixel rolling ball radius. Then, the threshold was automatically adjusted for the dark background with the default setting. By particle analysis, the fluorescence cells were fit to ellipses, and their major and minor length was measured as cell length and width, respectively, with cut off by more than 1 $\mu m^2$ and more than 0.7 circularity of the particles.

*Size and Concentration Measurement of Purified Extracellular Vesicles by Nanoparticle Tracking Analysis.* The purified EVs were diluted in 0.2-μm pore filtered DPBS to about $10^7$–$10^8$ orders particles/mL and analyzed using NanoSight NS300 (NanoSight, Amesbury, UK) with a 488-nm blue laser, a clear filter, and NTA 3.4 Built 3.4.4 software at a shutter speed of 1500, camera gain of 500, and threshold of five for 60 s at 20 ºC five times for each sample.

### Supporting Figures

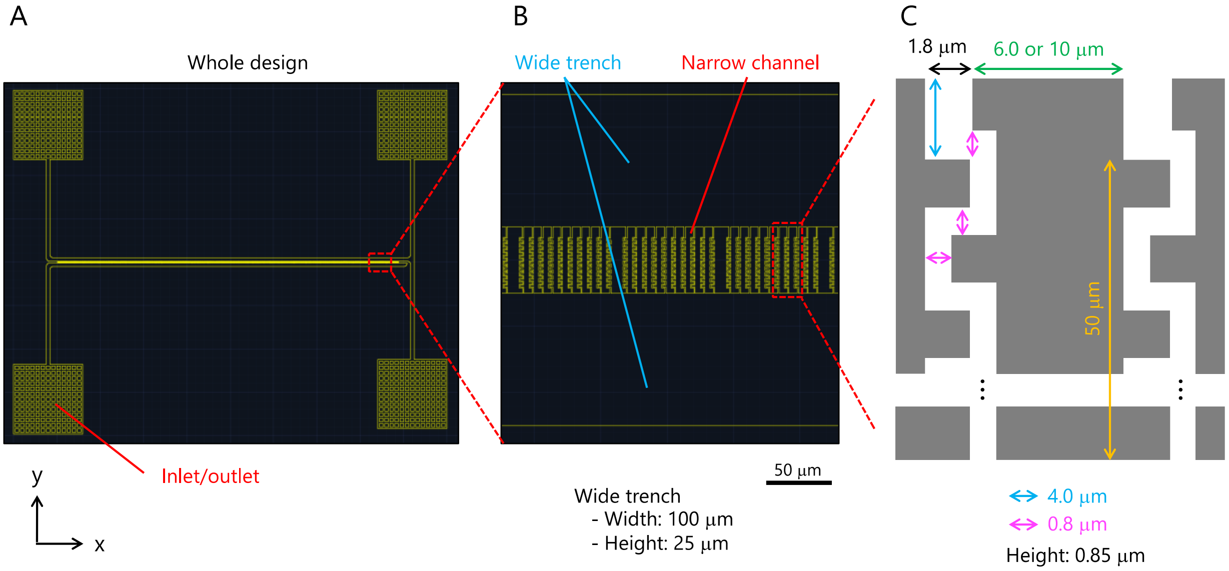


**Figure S1.** Design of microfluidic device and narrow channels with winding structures. (A) The whole design of the iMM microfluidic device from the top view. The device has four inlets/outlets. A wide trench connects three inlets/outlets at the upper left, upper right, and lower right. Narrow channels connect this wide trench with the other one from the lower-left inlet/outlet. (B) A magnified image of narrow channels. Narrow channels with winding traps connect two wide trenches. (C) A sketch of narrow channels with a winding trap. Cells are loaded by the flow running the wide trench at the top of the sketch and trapped at a cell trap, indicated by a blue arrow. The sketch is not to scale.

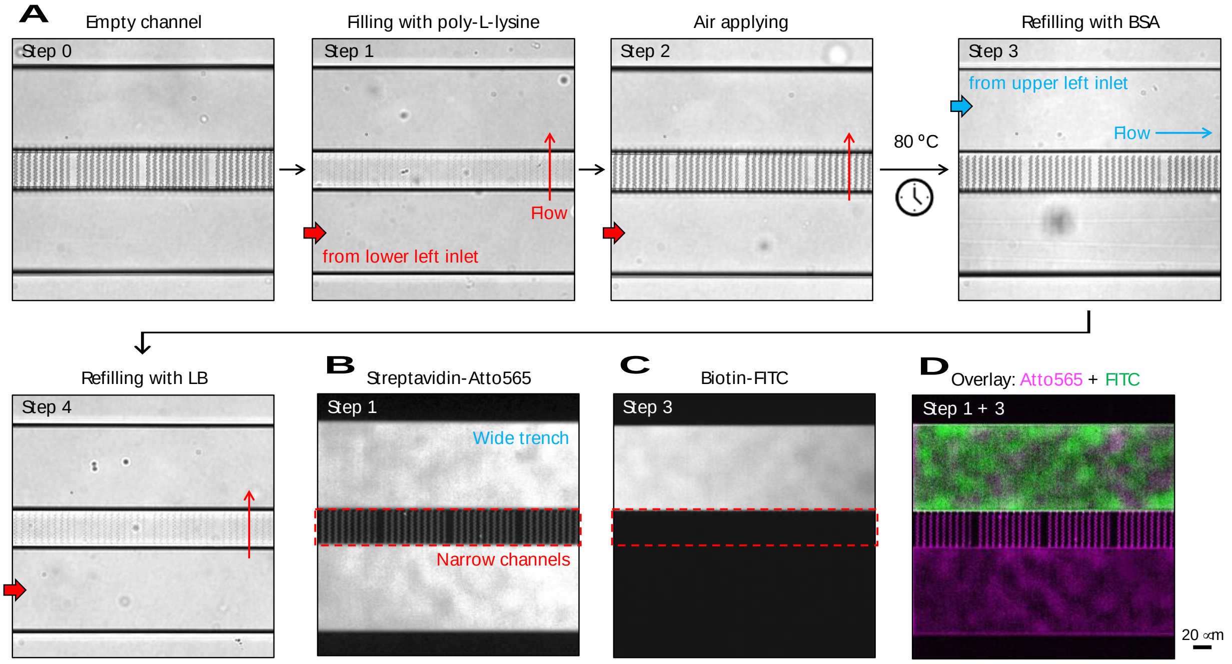


**Figure S2.** Surface modification of the microfluidic device. (A) Brightfield images of the iMM microfluidic device during the process of surface patterning. (B–D) Fluorescence images for illustration, (B) the device filled with streptavidin-Atto565 instead of poly-L-lysine at Step 1. (C) the device filled with biotin-FITC instead of poly-L-lysine at Step 3. (D) Overlayed and colored images of B and C. Green: biotin-FITC, and magenta: streptavidin-Atto565.

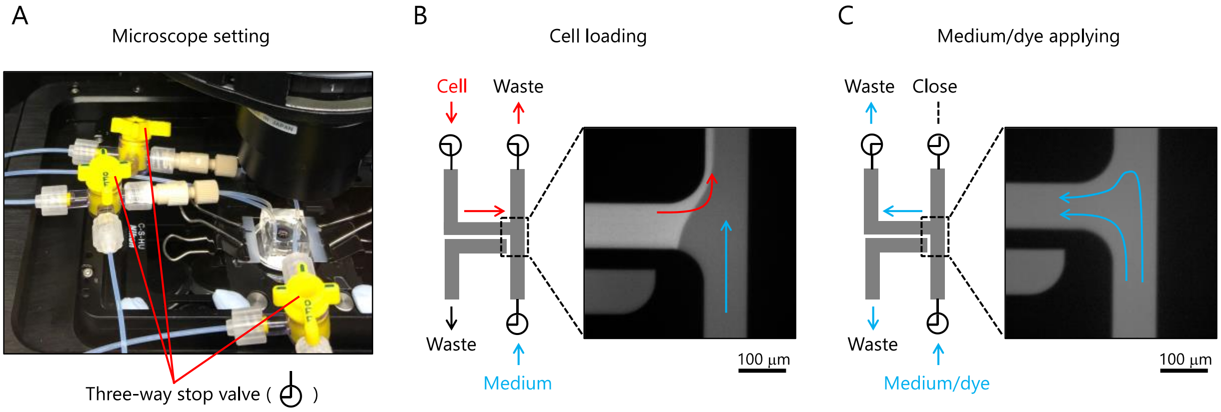


**Figure S3.** Prevention of contamination in a medium inlet by flow control using three-way stop valves. (A) An image of microscope settings. Three three-way stop valves were connected to the inlets/outlets of the microfluidic device via tubing. (B) Flow direction in the device for cell loading. A sketch of the device with inlets/outlets is on the left, and a representative fluorescence image of the device at the cell loading step is on the right. Cells and medium were applied from the upper-left and the lower-right inlets, respectively. The medium applied at the cell loading step prevented contamination of the inlet. (C) Flow direction in the device for medium/dye applying. A sketch of the device with inlets/outlets is on the left, and a representative fluorescence image of the device at the medium applying step is on the right. The medium was applied from the lower-right inlet and to the upper-left outlet because the upper-left outlet was open to the waste bottle, and the upper-right outlet was closed. All images were taken by multiple independent experiments. The sketches are not to scale.

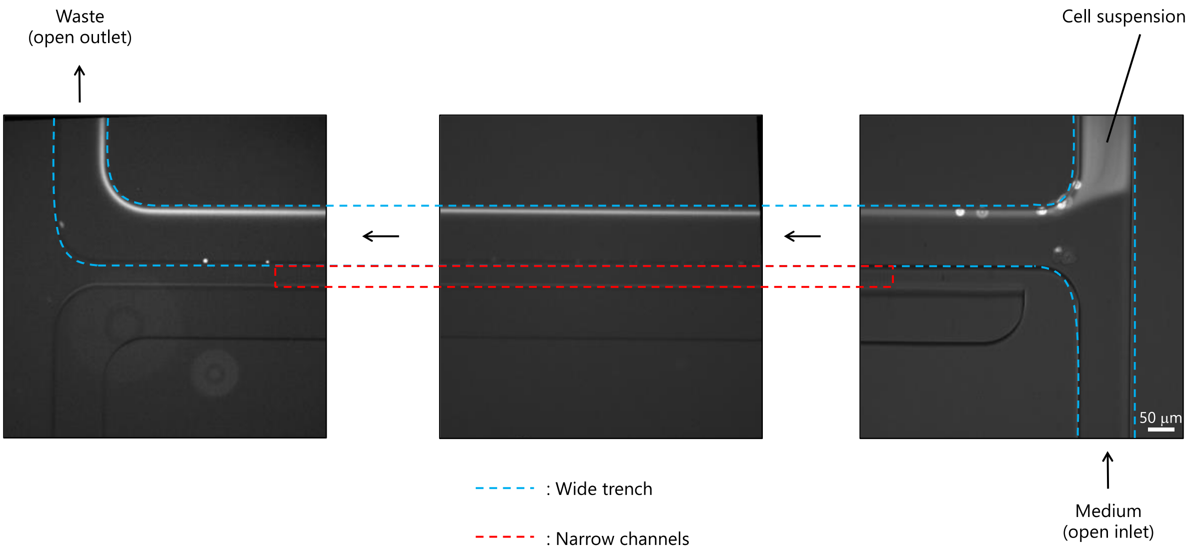


**Figure S4.** Prevention of contamination from narrow channels by laminar flow. Representative fluorescence images of iMM microfluidic device at three different positions are shown. First, the medium was applied from the lower-right inlet to the upper-left outlet. Then, the cell suspension (as shown in white) at the upper-right inlet was carried along the wall by a laminar flow of the medium to the upper-left outlet, preventing the contamination of narrow channels. All images were taken by multiple independent experiments.

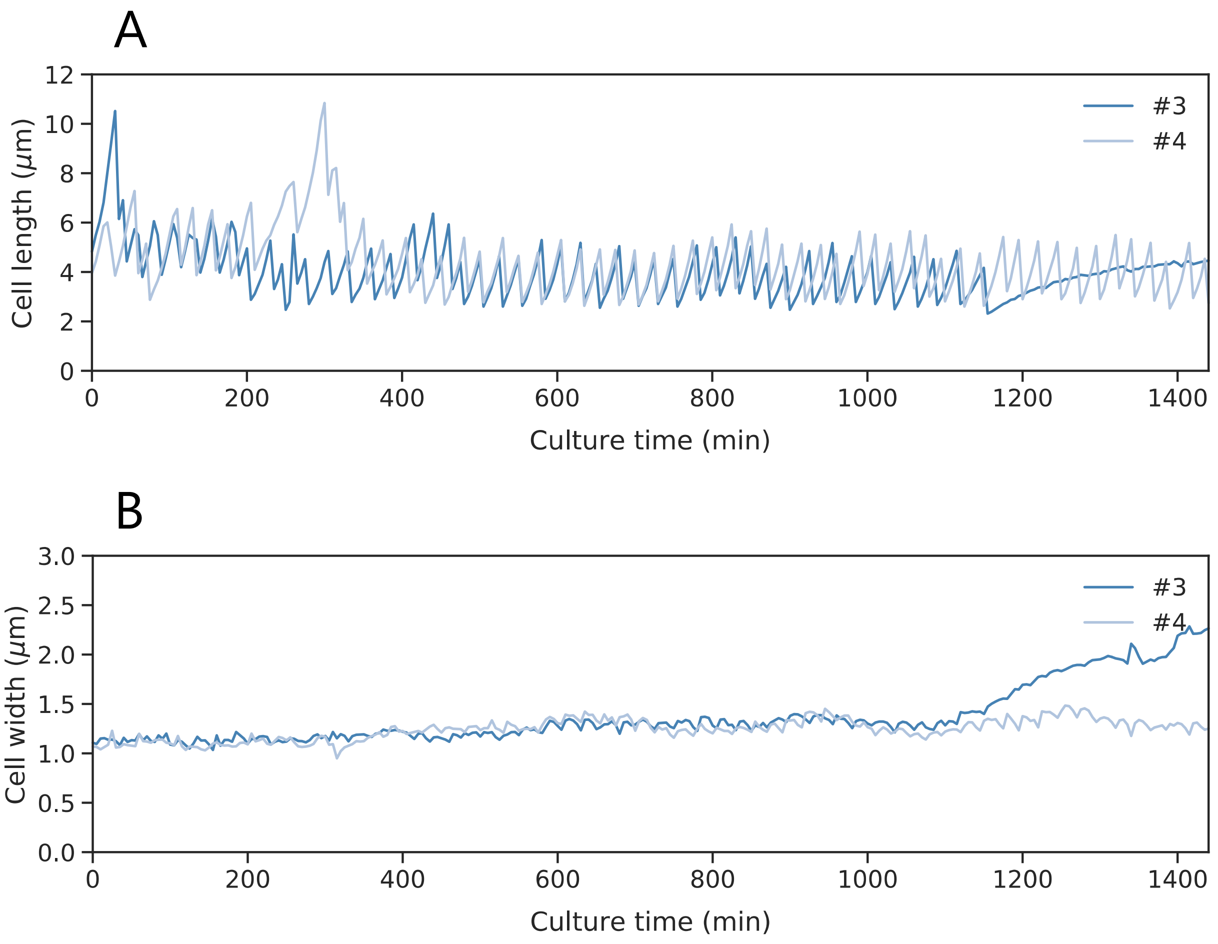


**Figure S5.** Additional data to Figure 2D and E showing the growth dynamics of captured *E. coli* cells. The growth of captured single cells was monitored for up to 24 h, and their cell length (A) and width (B) were measured from the fluorescence images. Each data line indicated single bacterial cells.

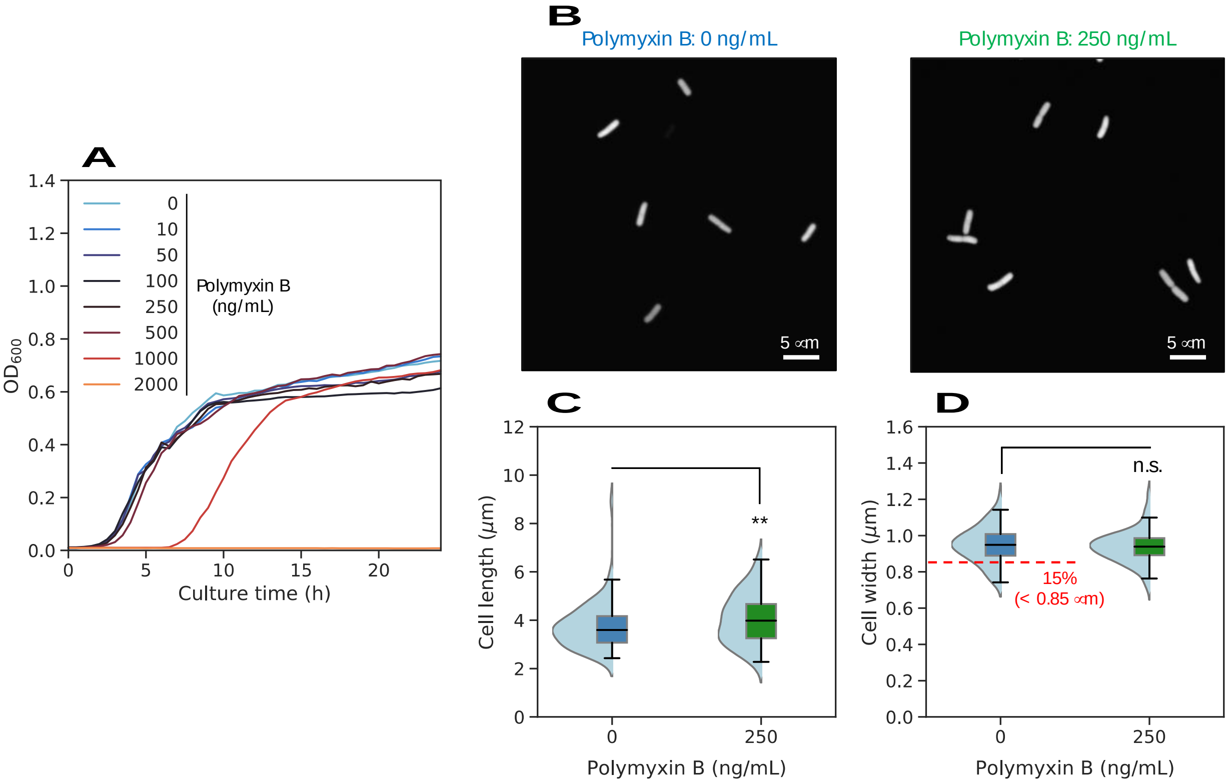


**Figure S6.** Growth and morphology of *E. coli* cells under low concentrations of polymyxin B in bulk. (A) Growth of *E. coli* cells at the different concentrations of polymyxin B. The lines and the error bands represent the mean and 95% confidence intervals of the data (n = 3), respectively. (B) Representative fluorescence images of GFP-expressing *E. coli* cells at the exponential growth phase and polymyxin B concentrations of 0 and 250 ng/mL (left and right, respectively). (C) Cell length obtained from the fluorescence images. (D) Cell width obtained from the fluorescence images. Approximate 15% of the cells were smaller in width, less than 0.85 μm, as indicated with a red dashed line. Each black line within the boxes represents the median, each lower and upper edges of the box represent the 25 and 75 percentiles of the data, respectively, and each error bar represents the maximum and minimum values of the box plots (n > 82). The gray dots on the right side of the boxes represent each data point. The light blue graphs on the left side of the boxes represent each data distribution. ***p* < 0.05 and n.s.: no significant difference (two-sided Brunner-Munzel test).

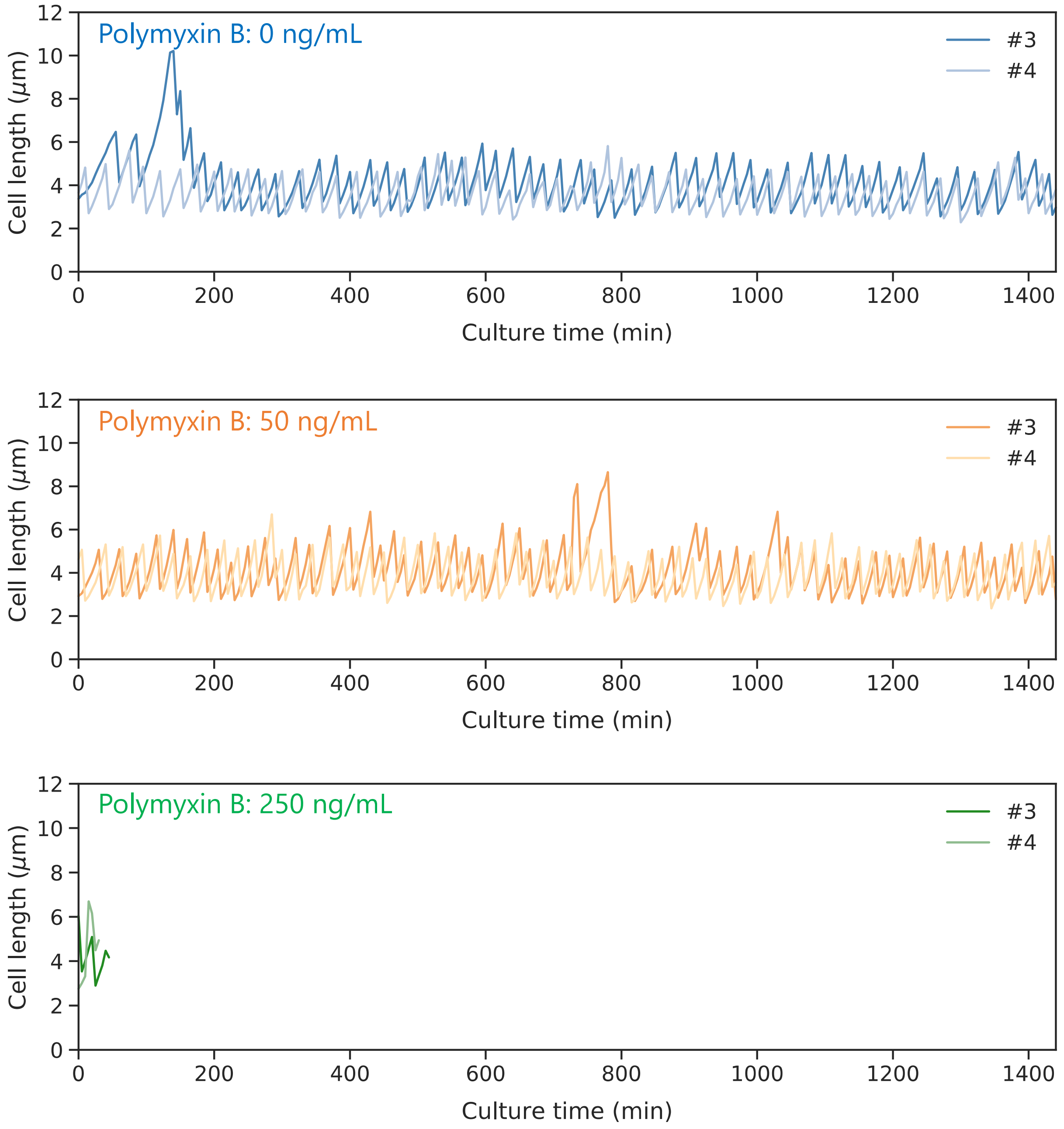


**Figure S7.** Additional data to Figure 3A, heterogeneous growth of single *E. coli* cells under antibiotic exposure. Cell length dynamics at different concentrations of polymyxin B, measured from fluorescence images. Each data line indicates single bacterial cells.

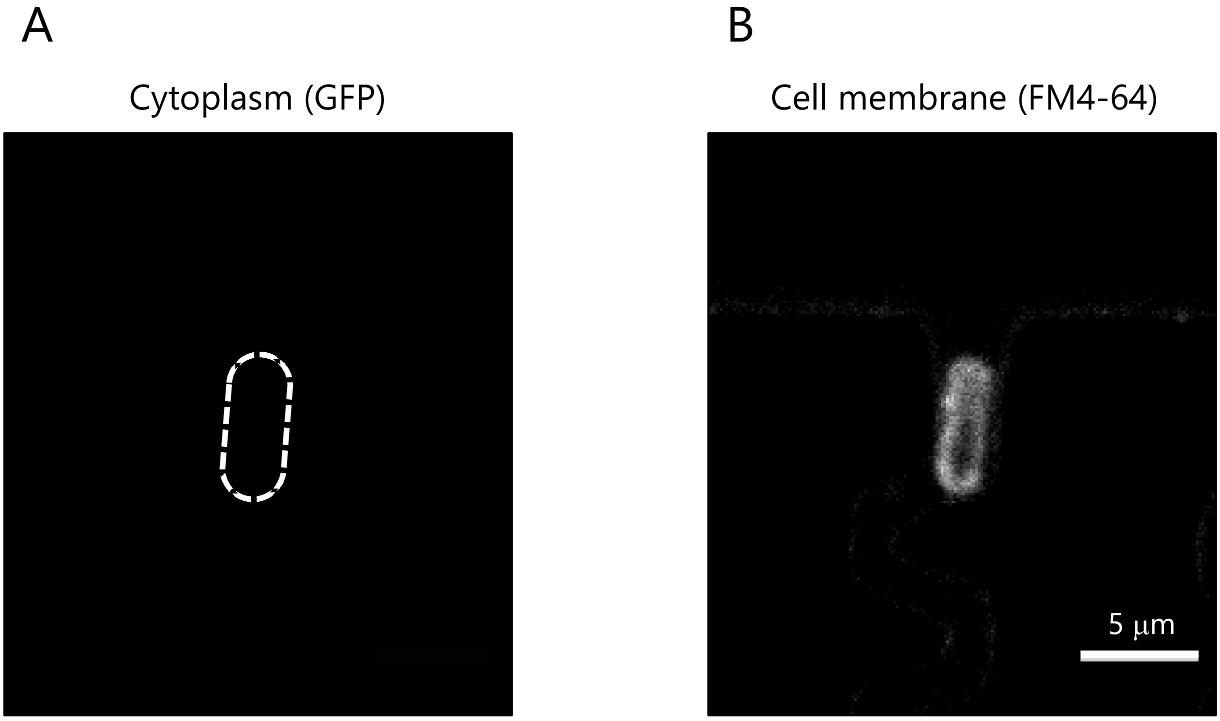


**Figure S8.** Dead cell harboring cellular membrane under polymyxin B. After 24-h culture under 250 ng/mL of polymyxin B and lipid staining with FM4-64, fluorescence images for GFP (A) and FM4-64 (B) were taken and set at the same brightness between them. The white dashed line in A shows the same position of the cell shown in B. The images without GFP fluorescence and with an FM4-64-labeled membrane mean that this cell died but the cell membrane is still visible (Cell #1 in Figure 3A bottom at polymyxin B 250 ng/mL). This cell death process is shown in Movie S5.

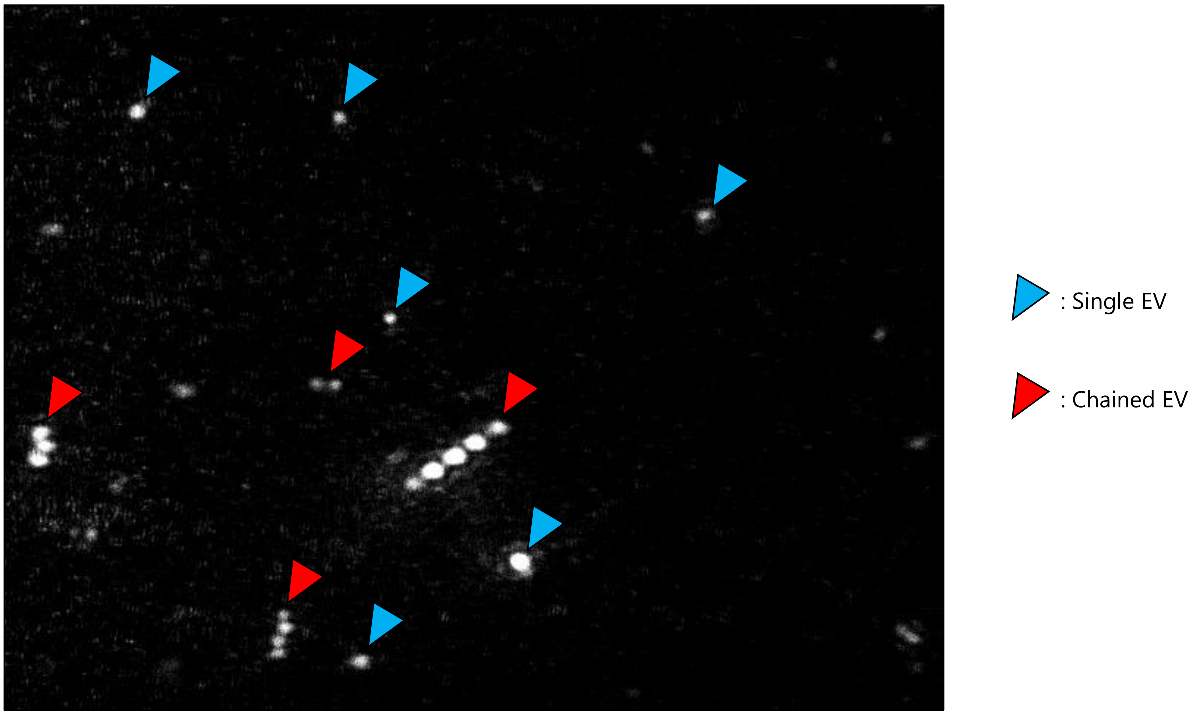


**Figure S9.** Morphology of single and chained EVs. An image of EVs taken from Movie S6 of nanoparticle tracking analysis is shown. These EVs were obtained from bulk culture supernatant without polymyxin B at the stationary growth phase. The median EV diameter was about 100 nm, according to Fig. S10B, by nanoparticle tracking analysis.

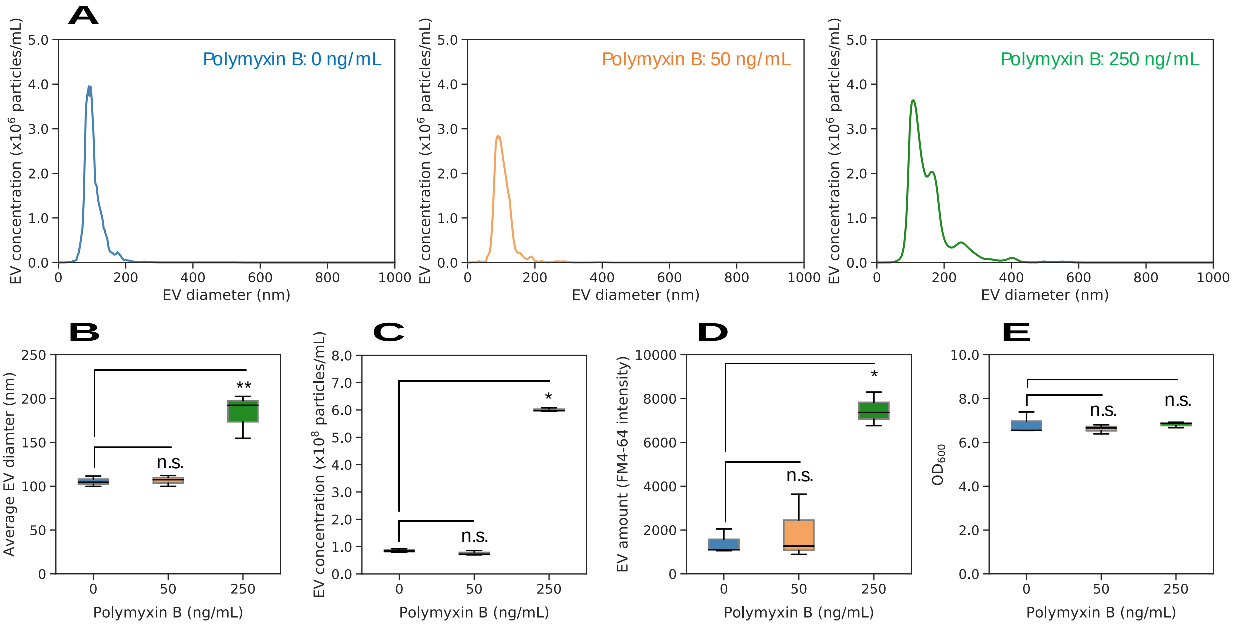


**Figure S10.** Induction of bigger EV secretion under low concentrations of polymyxin B. (A–C) Size distribution (A), average diameters (B), and average concentrations (C) of EVs at the stationary growth phase and polymyxin B concentration of 0, 50, and 250 ng/mL by nanoparticle tracking analysis. In A, the line and the error band represent the median and the 25 and 75 percentiles of the data (n = 3), respectively. (D) EV amounts at the stationary growth phase and different polymyxin B concentrations quantified with a lipid-staining dye, FM4-64. (E) $OD_{600}$ values at the stationary growth phase and different polymyxin B concentrations when the culture media were collected. Each black line within the boxes represents the median, each lower and upper edges of the boxes represent the 25 and 75 percentiles of the data, respectively, and each error bar represents the maximum and minimum values of the box plots (n = 3). The gray dots on the right side of the boxes represent each data point. *$p < 0.01$, **$p < 0.05$, and n.s.: no significant difference (two-sided Brunner-Munzel test).

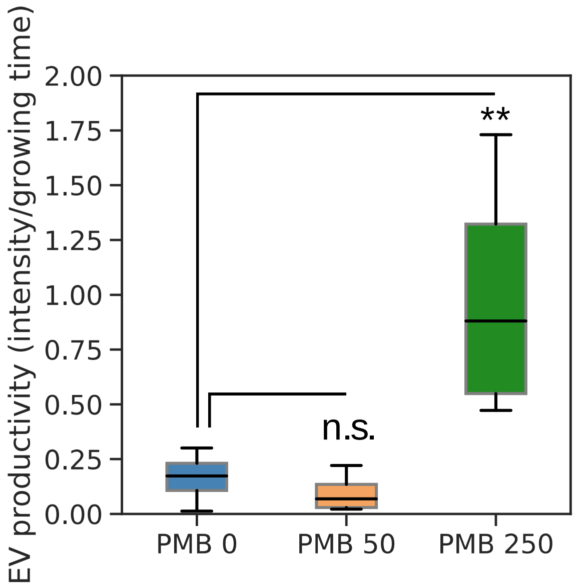


**Figure S11.** EV productivity of single mother cells at different concentrations of polymyxin B. The EV productivity was obtained by dividing the fluorescence intensity (Figure 5D) by the total growing time of each (Figure 3A and Figure S7). The EV productivity at 250 ng/mL of polymyxin B (0.88 a.u./min) was significantly higher, but that at 50 ng/mL of the antibiotics (0.07 a.u./min) was not more than that in the absence of polymyxin B (0.17 a.u./min). Each black line within the boxes represents the median, each lower and upper edges of the boxes represent the 25 and 75 percentiles of the data, respectively, and each error bar represents the maximum and minimum values of the box plots (n = 4). The gray dots on the right side of the boxes represent each data point. ***$p$ < 0.05, and n.s.: no significant difference (two-sided Brunner-Munzel test).

**Descriptions of the supporting movies.**

**Movie S1.** Growth dynamics of GFP-expressing *E. coli* cell without polymyxin B for 24 h (Cell #1 in Figure 2D and E). Fluorescence images of a GFP-expressing cell and brightfield images of a narrow channel were taken every 5 min for up to 24 h, were composited, and are shown at 13 frames per sec. Green: GFP-expressing *E. coli* cell.

**Movie S2.** Growth dynamics of GFP-expressing *E. coli* cell without polymyxin B for 24 h, followed by dividing fault (Cell #3 in Figure S5). Fluorescence images of a GFP-expressing cell of a narrow channel were taken every 5 min for up to 24 h and are shown at 13 frames per sec. The cell stopped the division at around 1160 min.

**Movie S3.** Growth dynamics of elongated GFP-expressing *E. coli* cell without polymyxin B for 24 h (Cell #1 in Figure 3A top). Fluorescence images of a GFP-expressing cell of a narrow channel were taken every 5 min for up to 24 h and are shown at 13 frames per sec. The cell elongated at around 565 min.

**Movie S4.** Growth dynamics of GFP-expressing *E. coli* cell with 50 ng/mL of polymyxin B for 24 h (Cell #1 in Figure 3A middle). Fluorescence images of a GFP-expressing cell of a narrow channel were taken every 5 min for up to 24 h and are shown at 13 frames per sec. The cell- derived fluorescence was getting faint and finally invisible at around 1200 min due to cell death by burst.

**Movie S5.** Growth dynamics of GFP-expressing *E. coli* cell with 250 ng/mL of polymyxin B for 24 h (Cell #1 in Figure 3A bottom). Fluorescence images of a GFP-expressing cell of a narrow channel were taken every 5 min for up to 24 h and are shown at 13 frames per sec. The cell-derived fluorescence was getting faint and finally invisible at around 90 min due to cell death.

**Movie S6.** EVs floating in a chamber used for nanoparticle tracking analysis. These EVs were obtained from culture supernatant without polymyxin B at the stationary growth phase. The median EV diameter was about 100 nm, according to Figure S10B, by nanoparticle tracking analysis.
